# Progranulin deficiency results in reduced bis(monoacylglycero)phosphate (BMP) levels and gangliosidosis

**DOI:** 10.1101/2021.09.30.461806

**Authors:** Sebastian Boland, Sharan Swarup, Yohannes A. Ambaw, Ruth C. Richards, Alexander W. Fischer, Shubham Singh, Geetika Aggarwal, Salvatore Spina, Alissa L. Nana, Lea T. Grinberg, William W. Seeley, Michal A. Surma, Christian Klose, Joao A. Paulo, Andrew D. Nguyen, J. Wade Harper, Tobias C. Walther, Robert V. Farese

## Abstract

Homozygous mutations of granulin precursor (*GRN*) lead to neuronal ceroid lipofuscinosis^1^, a severe neurodevelopmental disease, in humans and neuroinflammation in mice^2^. Haploinsufficiency of *GRN* almost invariably causes frontotemporal dementia (FTD)^3,4^. The *GRN* locus produces progranulin (PGRN), a lysosomal precursor protein that is cleaved to granulin peptides^5,6^. Despite intensive investigation, the function of granulins and the reason why their absence causes neurodegeneration remain unclear. Here, we investigated PGRN function in lipid degradation, a major function of lysosomes. We show that PGRN-knockout human cells, PGRN-deficient murine brain, and frontal lobes of human brains from patients with *GRN* mutation-related FTD have increased levels of gangliosides, highly abundant sialic acid–containing glycosphingolipids (GSL) that are degraded in lysosomes. Probing how PGRN deficiency causes these changes, we found normal levels and activities of enzymes that catabolize gangliosides. However, levels of bis(monoacylglycero)phosphate (BMP), a lysosomal lipid required for ganglioside catabolism^7^, were markedly reduced in PGRN-deficient cells and patient brain tissues. These data indicate that granulins are required to maintain BMP levels, which regulate ganglioside catabolism, and that PGRN deficiency in lysosomes leads to gangliosidosis. This aberrant accumulation of gangliosides may contribute to neuroinflammation and neurodegeneration susceptibility.

## INTRODUCTION

About half of the human brain mass is composed of lipids^8^. Lysosomes are crucial in degrading and recycling cellular lipids. Accumulation of lipids and other macromolecules due to lysosomal dysfunction is linked to many neurodevelopmental and neurodegenerative diseases, broadly classified as lysosomal storage disorders. Granulins localize to lysosomes, prompting us to examine the hypothesis that they function in lysosomal lipid metabolism. Consistent with this notion, a previous lipidomic study showed that PGRN deficiency in humans or mice alters levels of brain triglycerides (TAG), sterol esters (SE), and phosphatidylserine (PS)^9^. However, this study did not examine ganglioside GSLs. Ganglioside catabolism occurs in the lysosome, and defects in the abundance or activity of different enzymes that catabolize gangliosides result in severe neurological diseases, such as Tay-Sachs or Sandhoff diseases^10,11^ (**Fig. 1a**).

**Figure 1.**
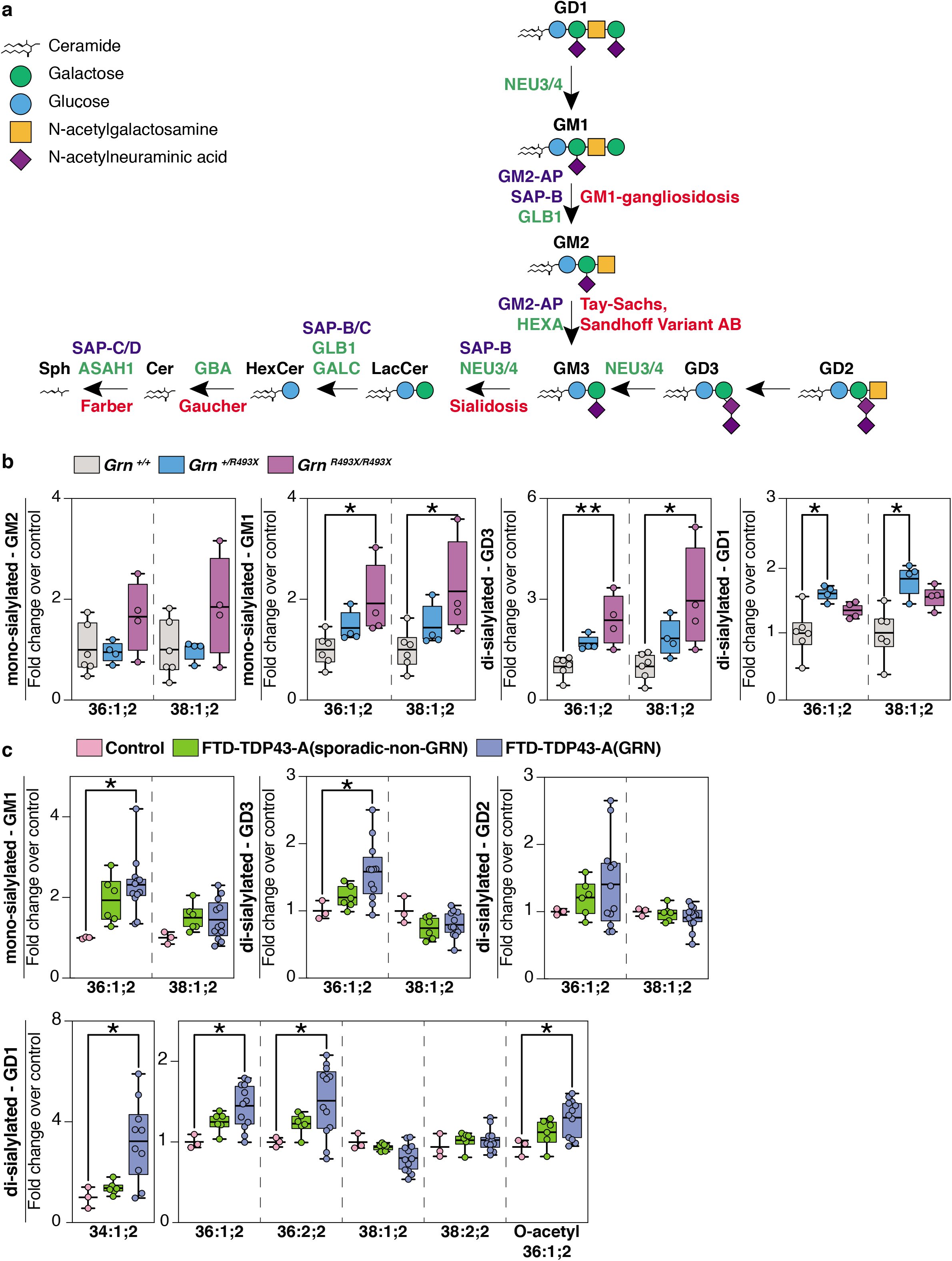
Deficiency of progranulin leads to ganglioside accumulation in mouse and human brain tissues. **a**, Ganglioside degradation pathway in the lysosome. The names of glycosyl hydrolases (green), activator proteins (purple), and associated metabolic diseases (red) are indicated in the scheme. **b**, Quantification of mono-sialyated and di-sialyated ganglioside species isolated from *Grn ^+/+^* (grey), *Grn ^+/R493X^* (blue), and *Grn ^R493X/R493X^* (purple) mouse brains. Box plots display mean ± the minimum and maximum number in the data set of six *Grn ^+/+^,* four *Grn ^+/R493X^* or four *Grn ^R493X/R493X^* biological replicates. **c**, Quantification of mono-sialyated and di-sialyated ganglioside species isolated from the frontal lobes of control (pink), FTD-TDP43-A (sporadic-non-GRN) (green), and FTD-TDP43-A (GRN) (blue) human brains. Box plots display mean ± the minimum and maximum number in the data set of three (control), six FTD-TDP43-A (sporadic-non-GRN) or 12 FTD-TDP43-A (GRN) biological replicates. GM2-AP, GM2 ganglioside activator protein; SAP-B/C/D, saposin-B/C/D; GLB1, galactosidase beta 1; HEXA, beta-hexosaminidase Subunit alpha; NEU3/4, neuraminidase 3/4; GALC, galactosylceramidase; GBA, glucosylceramidase beta; ASAH1, N-acylsphingosine amidohydrolase 1; SAP-C/D, saposin-C/D. One-way ANOVA followed by multigroup comparison (Dunn’s) test was performed. *p<0.05, **p<0.01, or ***p<0.001.

## RESULTS

We utilized lipidomics to examine GSLs in PGRN-deficient cells and tissues. We first analyzed brains isolated from 18-month-old *Grn ^R493X^* mice, a murine model of PGRN deficiency^12^. These mice harbor the murine equivalent of the most prevalent human *GRN* mutation that causes FTD (R493X), and they phenocopy *Grn* knockout mice, exhibiting CNS microgliosis, cytoplasmic TDP-43 accumulation, reduced synaptic density, lipofuscinosis, hyperinflammatory macrophages, and excessive grooming behavior^12^. Lipodomics revealed increased levels of mono-sialylated GM1 species and a two-to fourfold increase in di-sialylated GD3 species in brains of *Grn ^R493X/R493X^* mice. Levels of these gangliosides trended higher in *Grn ^+/R493X^* heterozygous mice, but did not reach significance. GM2 levels also trended higher in *Grn ^R493X/R493X^* mice, but were not significantly changed in *Grn ^R493X/+^* mice. Di-sialylated GD1 species were increased in brains of *Grn ^R493X/+^* brains and trended higher in *Grn ^R493X/R493X^* brains (**Fig. 1b and Extended Data Fig. 1a**). We also found modestly reduced levels of long-chain bases (LCB; sphingosine and sphinganine) in *Grn ^R493X/R493X^* brains, compared with control brains (**Extended Data Fig. 1a)**. Because sphingosines are generated by degradation of more complex sphingolipids^13^, their reduced levels suggest that degradation of sphingolipids is impaired in PGRN-deficient brains. Unlike sphingolipids, levels of the phospholipids phosphatidylethanolamine (PE), phosphatidylcholine (PC), PS, and of neutral lipids were similar among genotypes (**Extended Data Fig. 1a)**.

To test whether PGRN deficiency’s impact on the brain lipidome was also present in patients with GRN-related FTD, conserved in humans, we analyzed the lipid composition of post-mortem human frontal and occipital lobe brain tissue from control, sporadic FTD (sporadic FTLD-TDP, Type A), and *GRN* mutation-related FTD (GRN FTLD-TDP, Type A) subjects. As reported for PGRN-deficient fibroblasts and murine brain^9^, we found increased levels of SEs and TAGs in the frontal lobes of GRN FTLD-TDP subjects; in contrast, TAGs were unchanged and SEs were below the detection limit in the occipital lobes of the same subjects (**Extended Data Fig. 1b**). Additionally, we found reductions in PE and cardiolipins (CL) (**Extended Data Fig. 1b**) and increases in sphingomyelin (SM), particularly in the frontal lobes of the patients with GRN FTLD-TDP.

Human brains are particularly rich in a variety of gangliosides, including GM1, GD1a/b, GD3, and GT1b^14^. In a pattern that was similar to the changes in the murine brain, we detected increased levels of mono-sialylated GM1 and di-sialylated GD3 and GD1 species in GRN FTLD-TDP subjects (**Fig. 1c**). Some of these ganglioside species also trended higher in sporadic FTLD-TDP subjects. The abundance of GT1, which can be catabolized at the plasma membrane^15^, was lower in GRN FTLD-TDP subjects and unchanged in sporadic FTLD-TDP subjects (**Extended Data Fig. 1b**). In contrast to the findings in the frontal lobes, we did not detect increases in gangliosides in the occipital lobes of either FTLD group (**Extended Data Fig. 1b)**.

To address the mechanism of ganglioside accumulation, we established HeLa cell lines with PGRN deficiency and such cells with PGRN expression restored (*GRN^-/-^* + addback) via lentiviral infection with untagged human PGRN cDNA (**Fig. 2a**). In HeLa cells, the most abundant ganglioside class is GM2 and the gangliosides GM3, GD3, and GD1a^16^ are present in lesser amounts. Levels of GM2 were ~twofold greater in PGRN-knockout cells than control cells, and normal levels were restored by PGRN-addback (**Fig. 2b**). GD1 and GD3 levels were not different in PGRN-knockout cells, but GD3 levels were reduced in the PGRN-addback cells (**Fig. 2b**). TAG levels were increased in PGRN-knockout cells and restored by PGRN-addback, whereas PC, PE, and PS levels were unaffected upon PGRN depletion. For unclear reasons, diacylglycerol (DAG) levels (**Fig. 2c**) and several sphingolipid catabolic products were unchanged in the PGRN-knockout cells but were increased in the PGRN-addback cells (**Extended Data Fig. 2a**). In agreement with the increase in GM2 gangliosides detected by lipidomic analyses, immunostaining of PGRN-knockout cells with an antibody that detects GM2 showed more cells with detectable GM2 puncta and more GM2 puncta per cell than control cells (**Fig. 2d**). These GM2 puncta co-localized with the lysosomal marker LAMP1 (**Extended Data Fig. 2b**), and their accumulation was reversed by PGRN-addback (**Fig. 2d**).

**Figure 2.**
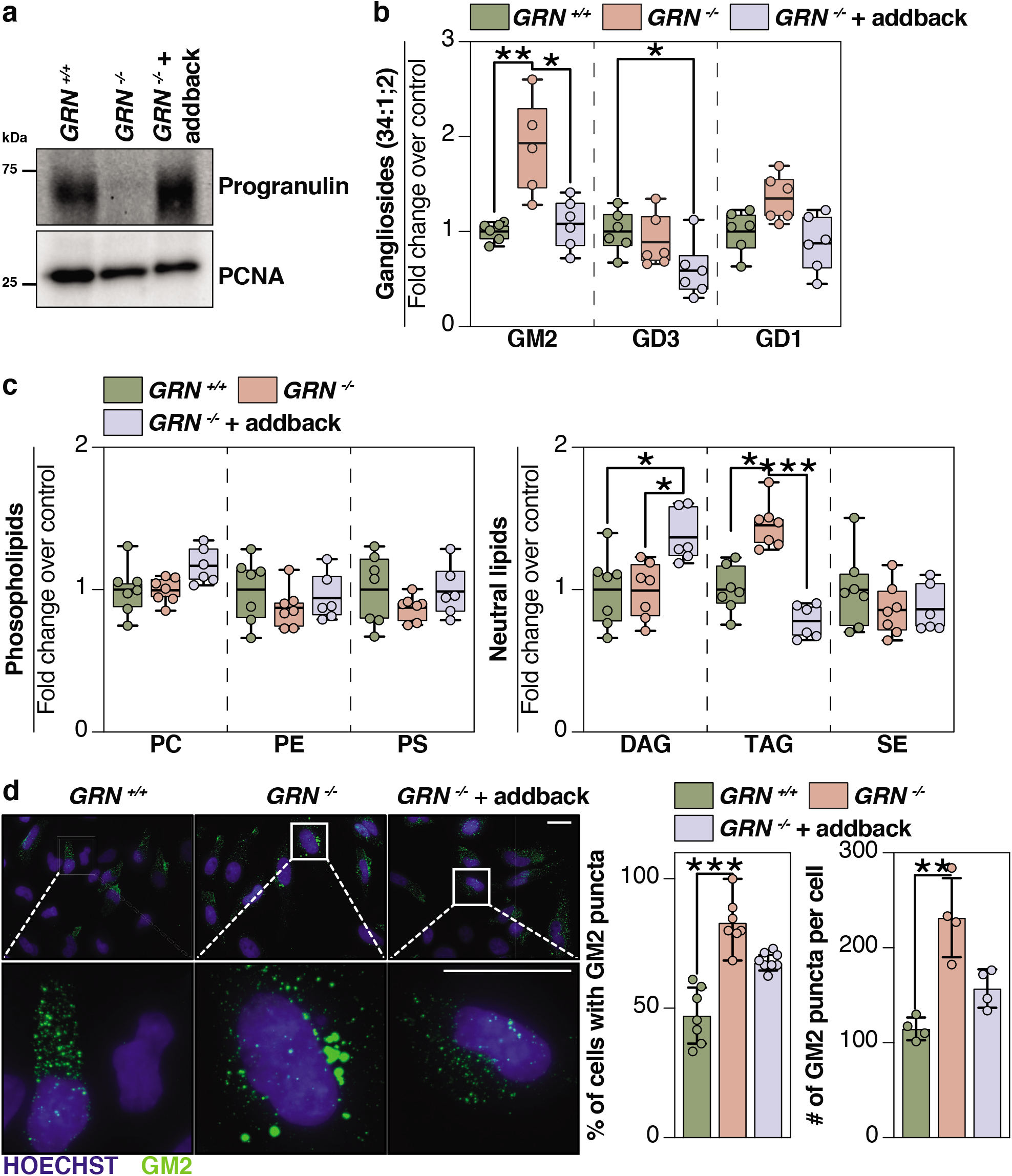
Lipidomic analysis and immunofluorescence analysis of HeLa cells reveals GM2 accumulation upon loss of progranulin that is restored by PGRN-addback. **a**, Western blot of full-length progranulin protein levels in *GRN^+/+^, GRN^-/-^,* and *GRN^-/-^* + addback HeLa cell lines. PCNA, proliferating cell nuclear antigen. **b**, Quantification of gangliosides and **c**, quantification of phospholipids (PC, PE, PS) and neutral lipids (DAG, TAG, SE) isolated from *GRN^+/+^* (green), *GRN^-/-^* (orange), and *GRN^-/-^* + addback (blue) HeLa cell lines. **d,** Representative confocal image of fixed HeLa cell genotypes stained with anti-GM2 antibody (green) and Hoechst (blue). Scale bar, 25 μm. Bar graphs display (left) percentage of cells positive for GM2 puncta and (right) number of GM2 puncta per cell. One-way ANOVA followed by multigroup comparison (Dunn’s) test was performed. *p<0.05, **p<0.01, or ***p<0.001. The number of cells used to calculate the percentage of GM2-positive cells were n=103, n=93, n=129 for *GRN^+/+^, GRN^-/-^,* and *GRN^-/-^* + addback, respectively. The number of cells used to calculate the GM2 puncta per cell were n=59, n=47, n=72 for *GRN^+/+^, GRN^-/-^,* and *GRN^-/-^* + addback, respectively. PC, phosphatidylcholine; PE, phosphatidylthanolamine; PS, phosphatidylserine; DAG, diacylglycerol; TAG, triacylglycerol; SE, sterol-esters.

Gangliosides are catabolized by lysosomal enzymes, and deficiency of these enzymes in abundance or activity leads to lysosomal lipid accumulation. We, therefore, tested lysosomal function in a number of assays. In quantitative whole-cell and lysosomal tandem mass tag (TMT) proteomics, we found no major differences in the abundances of lysosomal proteins or GSL-metabolizing enzymes in PGRN-knockout and PGRN-addback cells **(Fig. 3a-d and Extended Data Fig. 3a)**. Moreover, the activity of the glycosphingolipid catabolism enzyme β-hexosaminidase subunit a (HEXA) was unchanged in PGRN-knockout and PGRN-addback genotypes when incubated with artificial substrates (**Fig. 3e**). The activity of glucosylceramidase β (GBA) was more variable in *GRN^-/-^* cell lysates than in controls, and the average trended slightly lower (~15%). Similar to HeLa cells, mouse brains deficient for PGRN had most of the lysosomal proteome unaffected, but we found a modest upregulation of several enzymes that mediate GSL degradation. (**Extended Data Fig. 3b).** Furthermore, we found normal enzymatic activity of HEXA in murine and human brain protein lysates *in vitro.* Similar to cells, the activity of GBA measured in mouse brain lysates derived from *Grn ^R493X/+^* and *Grn ^R493X/R493X^* genotypes trended lower than in control brain lysates (**Extended Data Fig. 3c**). Notably, human brain lysates of the frontal lobe exhibited less activity of GBA when incubated with artificial substrates (**Extended Data Fig. 3c**). In contrast, brain lysates of the occipital lobe showed no differences in GBA activity (**Extended Data Fig. 3c**).

**Figure 3.**
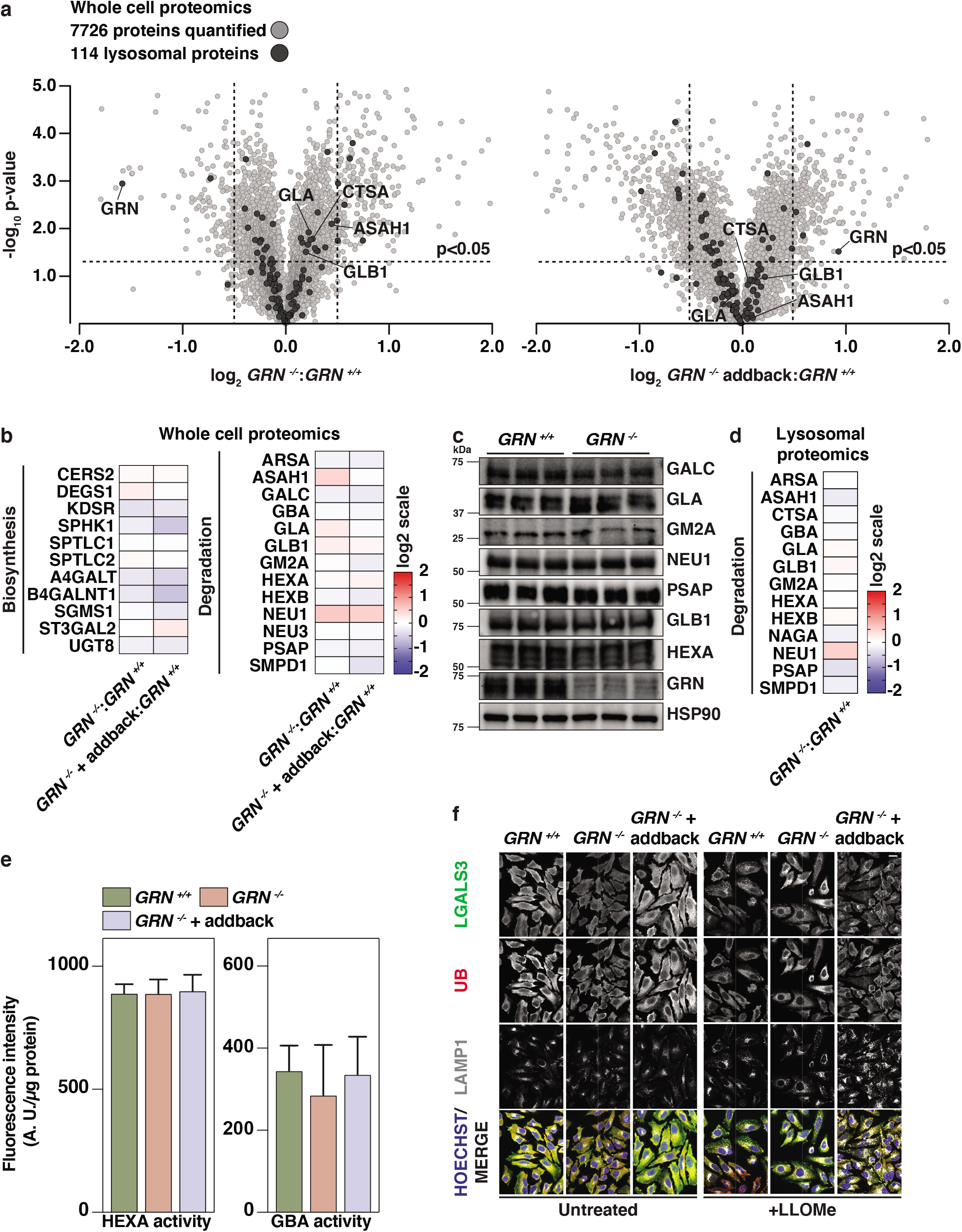
TMT-quantitative proteomic and in vitro analyses show no major differences in abundances or activities of glycosphingolipid metabolic enzymes in cells with PGRN depletion. **a**, Volcano plot representation of *GRN^-/-^* (left) and *GRN^-/-^* + addback (right) plotted against *GRN^+/+^* with log_2_-fold-change (ratio of relative abundance, x-axis) and -log_10_ p-value (y-axis). All proteins (grey) and lysosomal proteins (black) quantified are shown. A corrected p-value<0.05 was used to calculate statistically significant differences between genotypes. **b,** Heat map analysis from whole-cell lysates of the relative abundance of a subset of proteins that are involved in glycosphingolipid biosynthesis and degradation. **c**, Western blotting analysis of proteins that are involved in glycosphingolipid degradation. **d**, Heat-map analysis from lysosomal fractions of the relative abundance of a subset of proteins that are involved in glycosphingolipid degradation. **e**, HEXA and GBA activities were assessed in *GRN^+/+^* (green), *GRN^-/-^* (orange), and *GRN^-/-^* + addback (blue) cells when incubated with artificial substrates (four biological replicates in technical triplicates, mean ±SD). One-way ANOVA followed by multigroup comparison (Dunn’s) test was performed. *p<0.05, **p<0.01, or ***p<0.001. **f**, Representative confocal image of fixed HeLa cell genotypes stained with anti-LGALS3 antibody (green), anti-ubiquitin (red), anti-LAMP1 (white), and Hoechst (blue) in the absence or presence of L-leucyl-L-leucine methyl ester. Scale bar, 25 μm.

Upon screening for other lysosome-mediated processes, we found no differences for mTOR signaling, autophagic flux, or phosphorylation of MiT/TFE family of proteins in PGRN-deficient cells or tissues (**Extended Data Fig. 3d,e**). These results suggest that PGRN depletion has minimal effect on lysosomal composition and function under basal conditions. However, in response to the lysosome permeabilizing compound, L-leucyl-L-leucine methyl ester (LLOMe), significantly more of the membrane-damage sensing molecule, galectin 3 (LGALS3), was recruited to lysosomes in PGRN-knockout cells than controls (**Fig. 3f**). In addition, these cells exhibited more ubiquitin puncta staining on damaged lysosomes than controls, which was reversed by expression of full-length PGRN (**Fig. 3f**). These findings indicate that, although lysosomes appear mostly unchanged at baseline, they respond differently to a damaging insult.

Because the levels of the enzymes that catabolize gangliosides were not changed in PGRN-deficient cells or tissues, and because lysosomes appeared mostly intact and functioning, we searched for another cause for gangliosidosis. BMP plays a crucial role in GSL and ganglioside degradation in lysosomes^17^, and its levels are altered in many lysosomal storage diseases^18^. BMP is found in intralumenal lysosomal vesicles (ILVs) where its negatively charged phosphate headgroup may enable binding of lysosomal hydrolases^19,20^. We measured BMP levels in PGRN-knockout HeLa cells and found it ~50% reduced, whereas levels of the BMP isomer (and presumptive precursor) PG were unchanged (**Fig. 4a**). The reductions in BMP levels were restored by addback of PGRN (**Fig. 4a**). We also examined BMP accumulation in the HeLa cell model system with radioactive tracers. Metabolic labeling studies utilizing ^14^C-arachidonic acid confirmed the reduction of BMP levels in PGRN knockout HeLa cells (**Fig. 4b**), suggesting that alterations in the synthesis or degradation of BMP with polyunsaturated fatty acids were impaired in PGRN deficiency. Similarly, brains of PGRN-deficient mice also showed a 50–60% reduction in BMP levels, and all detected BMP species were significantly reduced in *Grn ^R493X/R493X^* brains (**Fig. 4c and Extended Data Fig. 4a)**.

**Figure 4.**
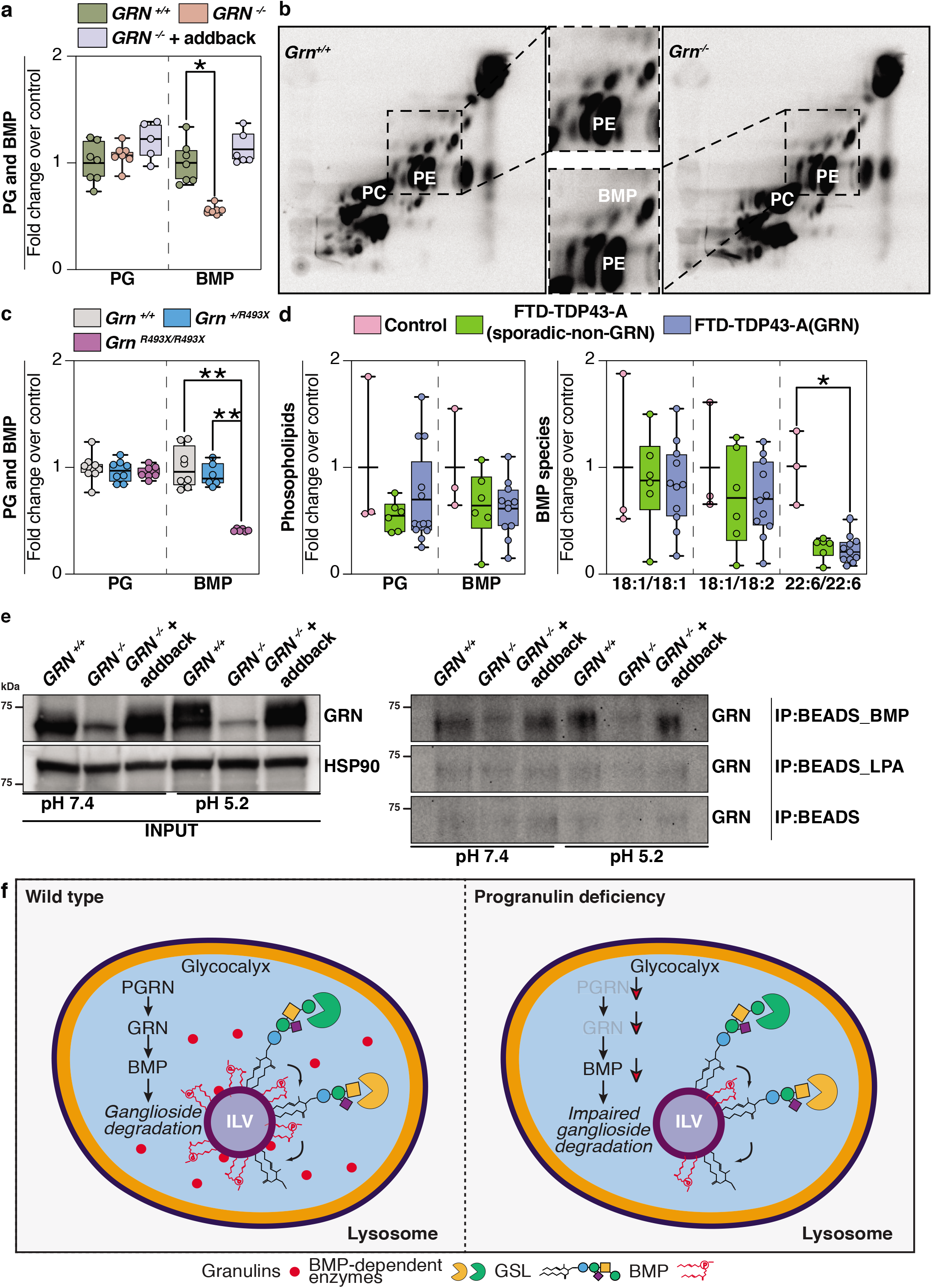
BMP levels are reduced in progranulin-deficient cells or brain tissues. **a**, Quantification of PG and BMP isolated from *GRN^+/+^* (green), *GRN^-/-^* (orange), and *GRN^-/-^* + addback (blue) HeLa cell lines reveals a ~50% reduction in BMP levels in PGRN-knockout HeLa. **b**, Labeling of cellular lipids by feeding ^14^C-arachidonic acid-albumin complex to *GRN^+/+^* and *GRN^-/-^* HeLa cell lines (60 minutes). Inlet highlights reduced levels of BMP in *Grn^-/-^* HeLa cells, compared to control cells. **c**, Quantification of PG and BMP isolated from *Grn ^+/+^* (grey), *Grn ^+/R493X^* (blue), and *Grn ^R493X/R493X^* (purple) mouse brains reveals a similar decrease in BMP levels upon loss of PGRN in mouse brains (~50%). **d**, Quantification of PG and BMP isolated from the frontal lobes of control (pink), FTD-TDP43-A(sporadic-non-GRN) (green), and FTD-TDP43-A(GRN) (blue) human brains. BMP species with mono- or di-unsaturated fatty acid moieties are not different, whereas BMP species containing two docosahexanoic acid moieties (22:6/22:6) are reduced in the frontal and occipital lobes of all FTD subjects. **e**, (left panel) Scheme for immunoprecipitation experiment of full-length PGRN using BMP-, LPA-coated, or control beads at pH 7.4 or pH 5.2 from *GRN^+/+^, GRN^-/-^,* and *GRN^-/-^* + addback cell lysates. (right panel) Western blotting anaylsis of full-length progranulin binding to individual beads reveals binding of progranulin to BMP-coated beads, in particular at pH 5.2. **f,** Model of the role of progranulin in the degradation of gangliosides. PGRN/granulin deficiency leads to reduced BMP levels through unclear mechanisms. Reduced BMP levels contribute to impaired ganglioside degradation. Eventually, this leads to lysosomal dysfunction and downstream consequences, including neuroinflammation and neurodegeneration. Oneway ANOVA followed by multigroup comparison (Dunn’s) test was performed. *p<0.05, **p<0.01, or ***p<0.001.

We also measured BMP levels in brain tissues from patients with GRN FTLD-TDP and compared them with sporadic FTLD-TDP and control subjects. We found marked decreases in a BMP species containing two docosahexanoic acid moieties (22:6/22:6) in the frontal and occipital lobes of all FTLD subjects (**Fig. 4d and Extended Data Fig. 4b)**. BMP with two 22:6 polyunsaturated fatty acids is one of the most abundant BMP species in human brain^21^. BMP species with mono- or di-unsaturated fatty acid moieties trended lower in some samples but were not overall different among the study groups.

We considered the hypothesis that PGRN (or granulins) may interact directly with BMP, perhaps to protect it from degradation. To test this hypothesis, we performed an immunoprecipitation assay with BMP-coated beads. Endogenous and over-expressed full-length PGRN from cell lysates specifically bound to BMP-coated beads, but not to uncoated-beads or beads coated with a control lipid, lysophosphosphatidic acid (LPA) (**Fig. 4e)**. The binding of PGRN to BMP appeared to be stronger in a lysosome-like acidic condition (pH 5.2) than at a neutral pH (pH 7.4).

Our experiments lead to a model for PGRN function in which lysosomal granulin peptides maintain lysosomal function and homeostasis, including the levels of BMP, which are crucial for ganglioside catabolism (**Fig. 4f**). In the setting of PGRN deficiency, reduced granulins lead to lysosomal dysfunction and reduced BMP levels on intralumenal vesicles (**Fig. 4f**). Binding of PGRN to BMP, recently reported independently^22^, suggests a model in which GRN binding enables BMP recycling from ILVs by shielding it from lysosomal degradation. Whatever the mechanism, low BMP levels impair ganglioside catabolism and result in gangliosidosis. The resultant gangliosidosis may also compromise certain lysosomal functions, such as response to membrane pertubation, and eventually lead to other hallmarks of the disease such as protein aggregation (e.g., TDP-43^23,24^). In agreement with an important role of lysosomal degradation in FTD, mutation in *CHMP2B,* a component of the ESCRT machinery of intralumenal vesicle formation, also leads to FTD^25^.

## DISCUSSION

Our results are consistent with several reports linking PGRN deficiency to altered GSL catabolism. Specifically, PGRN has been linked to HEXA function^1^, and two reports found that GBA activity is compromised in PGRN-deficient mice or neurons^26,27^. While we found that HEXA activity was not changed, GBA activity was decreased in the frontal lobes (but not the occipital lobes) of GRN FTLD-TDP patients. Finally, and of particular relevance to our findings of PGRN deficiency causing gangliosidosis, a recent report corroborates reduced BMP levels in brains of PGRN-deficient mice^22^.

Despite the widespread expression of PGRN in tissues, PGRN deficiency results primarily in a disease of the central nervous system. Our findings suggest that this may be because gangliosides are most prevalent in the CNS. In the brain, neuronal gangliosidosis may trigger the activation and recruitment of microglia to phagocytose and process the excess gangliosides. In particular, GM2 gangliosides may incite TNF-a expression and inflammation in monocyte-derived cells^28^. A hallmark of PGRN deficiency in murine brain is microgliosis and neuroinflammation^2^, and microglial cells (and macrophages) are hyperactivated in the setting of PGRN deficiency^2^. The accumulated effects of long-term defects in lysosomal ganglioside metabolism and neuroinflammation may, therefore, contribute to GRN FTLD-TDP.

With respect to therapeutic implications, increased ganglioside levels or reduced BMP levels may serve as biomarkers for PGRN-deficient FTLD or other neurodegenerative disorders. It may also be of interest to explore whether drugs that lower ganglioside production^29^ are beneficial in GRN-FTLD-TDP.

## ACKNOWLEGMENTS

We thank members of the Farese & Walther laboratory, Gilbert Di Paolo and Todd Logan (Denali Therapeutics) and Konrad Sandhoff for helpful discussions. We thank Gary Howard for editorial services. This work was supported by the Bluefield Project to Cure FTD (SB, TCW, and RVF), NIH grants R01NS083524, R01NS110395 (to JWH) and R01GM132129 (to JAP), the Harvard Brain Initiative ALS seed grant program (to JWH), a generous gift from Ned Goodnow (JWH), the Canadian Institutes for Health Research (to S.S.), and the Howard Hughes Medical Institute (TCW). The UCSF Neurodegenerative Disease Brain Bank receives funding support from NIH grants P30AG062422, P01AG019724, U01AG057195, and U19AG063911, as well as the Rainwater Charitable Foundation and the Bluefield Project to Cure FTD.

## AUTHOR CONTRIBUTIONS

S.B., R.V.F. and T.C.W. designed the study; S.B., S.S., A.D.N., G.A., R.C.R. performed experiments; S.B. performed and analyzed lipidomics mass spectrometry experiments; Y.A.A., A.W.F., S.S. contributed with sample preparations; A.N.L., S.S., L.T.G, W.W.S. procured brain bank samples for biochemical analysis; S.B., M.S., C.K. performed and analyzed lipid mass spectrometry experiments of human brain samples; J.W.H. co-supervised cultured cell and proteomics experiments by S.S., S.B.; J.A.P provided mass spectrometry expertise for the proteomics experiments; S.B., S.S. created figures and S.B., S.S., R.V.F. and T.C.W. wrote the manuscript with comments from all authors.

---

**Extended Data Figure 1.**
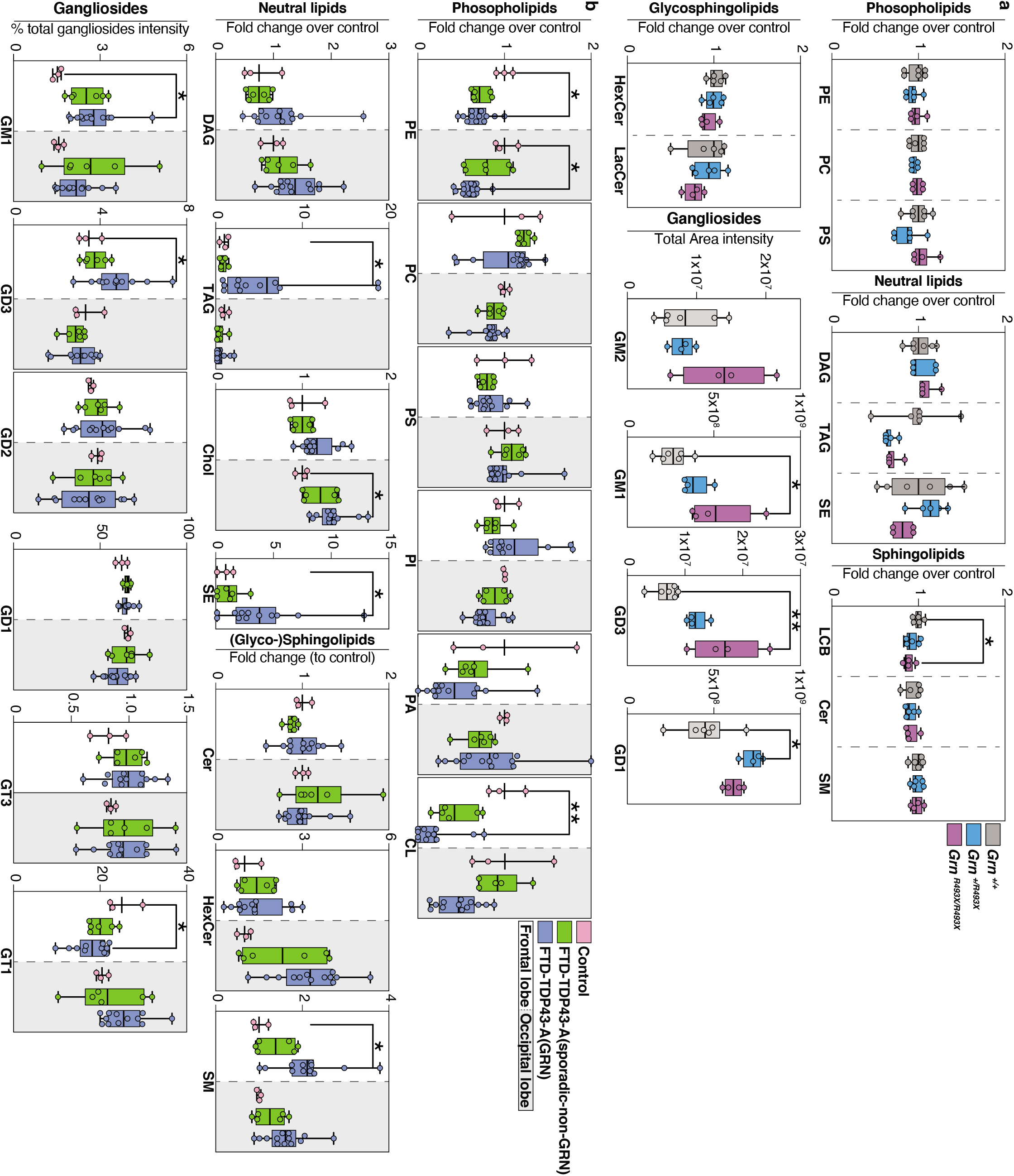
Global lipid analysis of mouse and human brains. **a,** Quantification of phospholipids (PE, PC, PS), neutral lipids (DAG, TAG, SE). (glyco-) sphingolipids (LCB, Cer, SM, HexCer, LacCer), and gangliosides (GM2, GM1, GD3, GD1) isolated from six *Grn ^+/+^* (grey), five *Grn ^+/R493X^* (blue), and four *Grn ^R493X/R493X^* (purple) biological replicates. PE, phosphatidylthanolamine; PC, phosphatidylcholine; PS, phosphatidylserine; DAG, diacylglycerol; TAG, triacylglycerol; SE, sterol-esters; LCB, long-chain base; Cer, ceramide; HexCer, hexosylceramide; LacCer, lactosylceramide; SM, sphingomyelin. P-values were calculated using one-way ANOVA followed by multigroup comparison (Dunn’s). *P<0.05. **b**, Quantification of phospholipids (PE, PC, PS, PI, PA, CL), neutral lipids (DAG, TAG, Chol, SE), (glyco-)sphingolipids (Cer, HexCer, SM), and gangliosides (GM1, GD3, GD2, GD1, GT3, GT1) isolated from the frontal and occipital lobes of control (pink), FTD-TDP43-A(sporadic-non-GRN) (green), and FTD-TDP43-A(GRN) (blue) human brains. Box plots display mean ± the minimum and maximum number in the data set. Data derived from the frontal lobes are shown in a box with a white background, and data derived from frontal lobes are shown in a box with grey shaded background. PE, phosphatidylthanolamine; PC, phosphatidylcholine; PS, phosphatidylserine; PI = phosphatidylinositol; PA, phosphatidic acid; CL, cardiolipin; DAG, diacylglycerol; TAG, triacylglycerol; Chol, cholesterol; SE, sterol-esters; Cer, ceramide; SM, sphingomyelin, HexCer, hexosylceramide; LacCer, lactosylceramide. One-way ANOVA followed by multigroup comparison (Dunn’s) test was performed. *p<0.05, **p<0.01, or ***p<0.001.

**Extended Data Figure 2.**
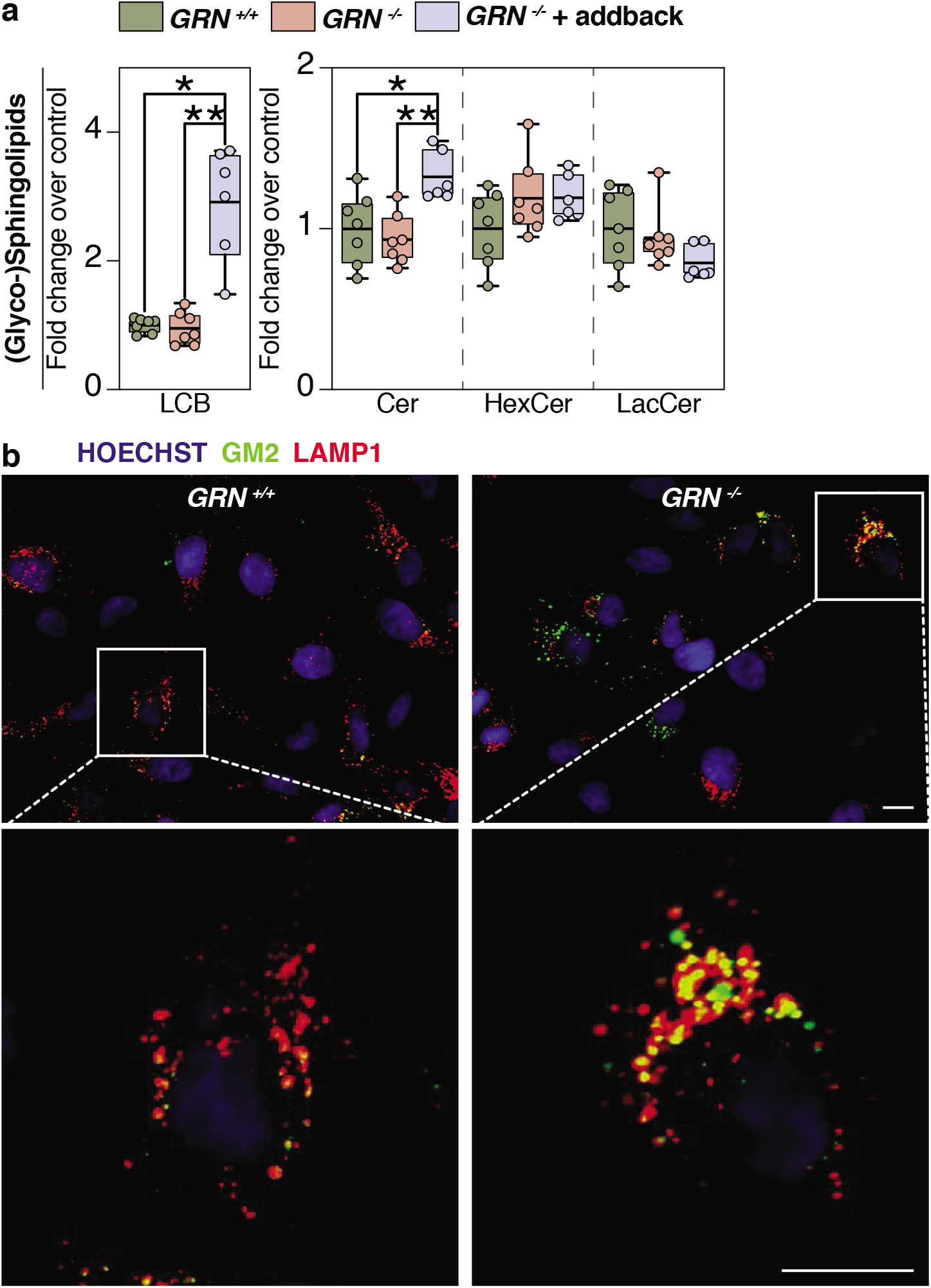
Loss of progranulin leads to GM2 accumulation in the lysosomes. **a,** Quantification of (glyco-)sphingoliopids (Cer, HexCer, LacCer) isolated from *GRN^+/+^* (green), *GRN^-/-^* (orange), and *GRN^-/-^* + addback (blue) HeLa cell lines. **b**, Representative immunofluorescent confocal image of fixed HeLa cell genotypes stained with anti-GM2 antibody (green), anti-LAMP1 antibody (red) and Hoechst (blue). Scale bar, 25 μm. Box plots display mean ± the minimum and maximum number in the data sets. LCB, long-chain base; Cer, ceramide; HexCer, hexosylceramide; LacCer, lactosylceramide. One-way ANOVA followed by multigroup comparison (Dunn’s) test was performed. *p<0.05, **p<0.01, or ***p<0.001.

**Extended Data Figure 3.**
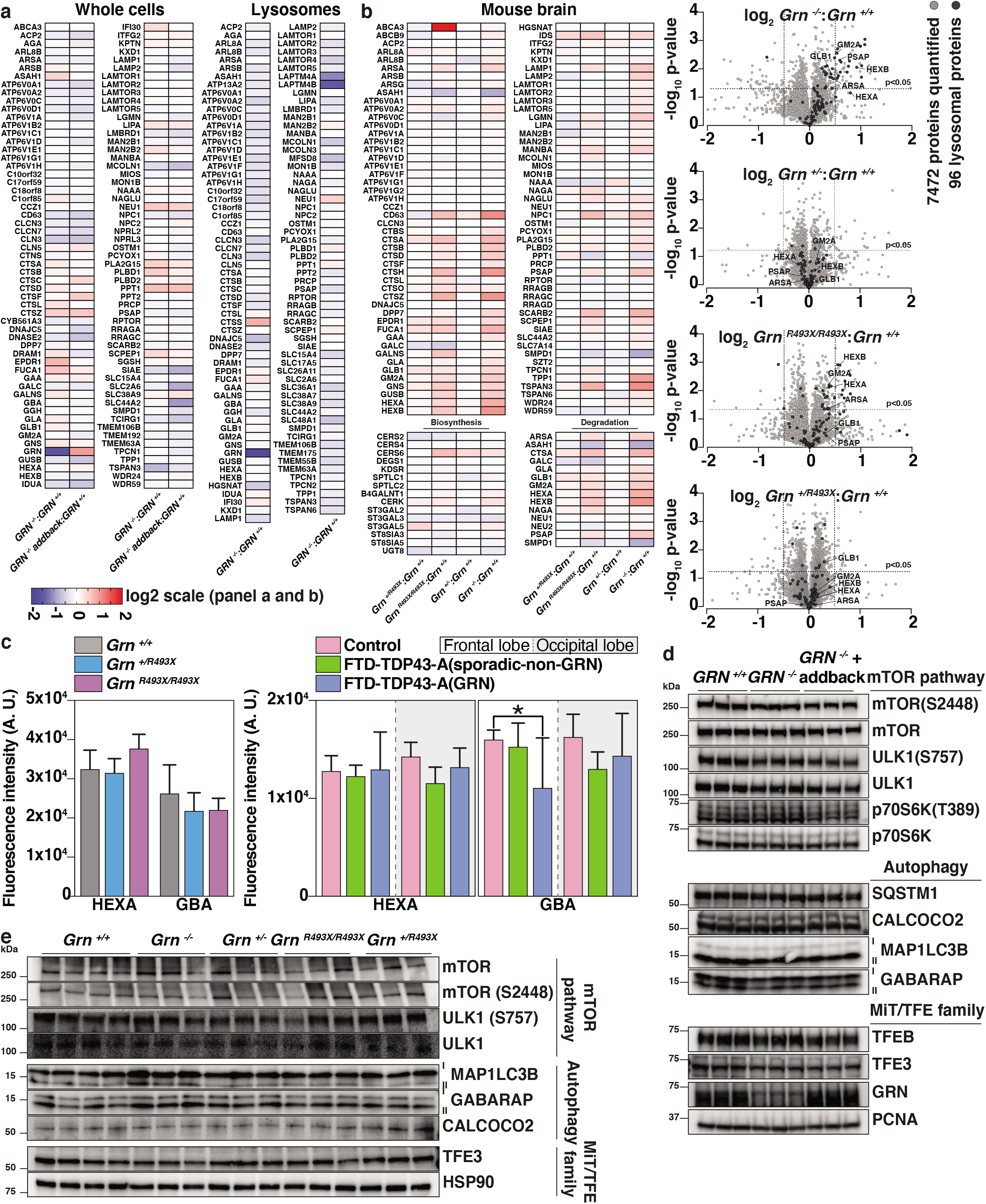
Quantitative TMT-proteomics, in vitro assays, and lysosomal functional assays of HeLa cell and mouse brains do not show any strong perturbations upon PGRN depletion. **a,** Heat-map representation of the relative abundance of lysosomal proteins from HeLa whole-cell or lysosomal extracts (three replicates each) using the genotypes, *GRN^+/+^, GRN^-/-^,* and *GRN^-/-^* addback. **b,** Heatmap and volcano plot representation of the relative abundance of lysosomal proteins and glycosphingolipid metabolic enzymes from mouse brains (n=3 or 4 each) of the following genotypes: *Grn ^+/+^, Grn ^-/-^, Grn ^+/-^, Grn ^+/R493X^* and *Grn ^R493X/R493X^.* For the volcano plots, log_2_ fold-change (ratio of relative abundance) is on the x-axis and -log_10_ p-value on the y-axis. All proteins (grey) and lysosomal proteins (black) quantified are shown. A corrected p-value<0.05 was used to calculate statistically significant differences between genotypes. **c**, HEXA and GBA activities were assessed in *Grn ^+/+^, Grn ^+/R493X^* and *Grn ^R493X/R493X^* mouse brain lysates (left) and in the frontal and occipital lobes of control (pink), FTD-TDP43-A(sporadic-non-GRN) (green), and FTD-TDP43-A(GRN) (blue) of human brain lysates as indicated using artifical substrates (n = three biological and three technical replicates, ±SD). One-way ANOVA followed by multigroup comparison (Dunn’s) test was performed. *p<0.05, **p<0.01, or ***p<0.001. **d** and **e**, Western blotting analysis of phosphorylation of MiT/TFE family proteins (as measured by gel-shift assay of TFEB, TFE3), mTOR pathway activity (as measured by the phosphorylation of mTOR kinase and its substrates ULK1, p70S6K), and autophagic flux (as measured by lipidation of MAP1LC3B, GABARAP and levels of autophagy receptors SQSTM1, CALCOCO2) in (**d**) HeLa whole cell extracts (*GRN^+/+^, GRN^-/-^ GRN^-/-^* + addback) and (**e**) mouse brains (*Grn^+/+^, Grn^-/-^, Grn^+/-^, Grn^+/R493X^,Grn^R493X/R493X^).*

**Extended Data Figure 4.**
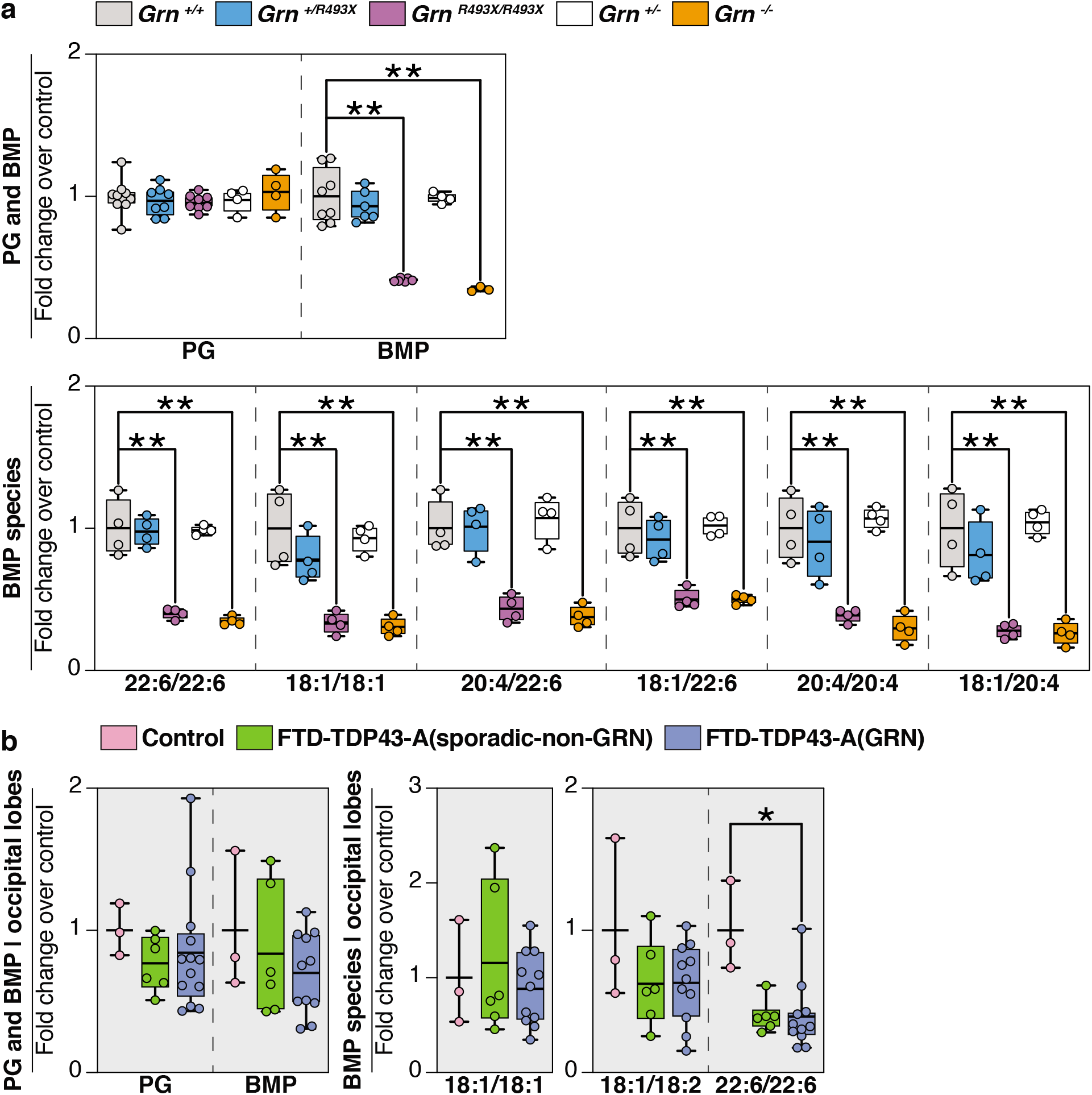
Loss of progranulin leads to reduced levels of BMP in cells, mouse brains, and human brains. **a,** Quantification of PG and BMP levels and individual BMP species from *Grn ^+/+^* (grey), *Grn ^+/R493X^* (blue), *Grn ^R493X/R493X^* (purple), *Grn ^+/+^* (white), and *Grn ^+/+^* (orange) mouse brains. **b,** Quantification of PG and BMP levels and individual BMP species from the occipital lobes of control (pink), FTD-TDP43-A(sporadic-non-GRN) (green), and FTD-TDP43-A(GRN) (blue) human brains. Box plots display mean ± the minimum and maximum number in the data sets. One-way ANOVA followed by multigroup comparison (Dunn’s) test was performed. *p<0.05, **p<0.01, or ***p<0.001.

## METHODS

### Chemicals and antibodies

The following reagents were purchased from commercial vendors: acetonitrile, methanol, water (all HPLC/MS grade), chloroform (HPLC grade, stabilized by 0.5-1% ethanol), ammonium formate, ammonium acetate, formic acid, and acetic acid were purchased from Sigma-Aldrich. N-Omega-CD3-octdecanoyl GM1 (Matreya LLC, Cat#2050), SPLASH® LIPIDOMIX® Mass Spec Standard ( Avanti Polar Lipids, Cat# 330707-1EA). TMTpro was purchased from Thermo Fisher (Cat#A44520). Primary antibodies against the following targets were used in the present study: GM2 monoclonal antibody (TCI America, Cat#A2576), human progranulin (R&D systems, Cat#AF2420), an anti-mouse progranulin polyclonal antibody that recognizes an epitope at amino acids 198–214^30^, PCNA (Santa Cruz Biotechnology Cat#sc56), GALC (Proteintech Cat#11991-1-AP), GLA (Proteintech Cat#66121-1-Ig), GM2A (Proteintech Cat#10864-2-AP), NEU1 (Santa Cruz Biotechnology Cat#sc166824), PSAP (Proteintech Cat#10801-1-AP), GLB1 (Proteintech Cat#15518-1-AP), HEXA (Proteintech Cat#11317-1-AP), HSP90 (Proteintech Cat#60318-1-Ig), TFEB (Cell Signaling Technology Cat#4240), TFE3 (Proteintech Cat#14480-1-AP), SQSTM1 (Proteintech Cat#18420-1-AP), CALCOCO2 (Proteintech Cat#12229-1-AP), MAP1LC3B (Cell Signaling Technology Cat#2775), GABARAP (Proteintech Cat#18723-1-AP), MTOR (Cell Signaling Technology Cat#2983), MTOR (S2448) (Cell Signaling Technology Cat#2971), P70S6K (Cell Signaling Technology Cat#2708), P70S6K (T389) (Cell Signaling Technology Cat#9234), ULK1 (Cell Signaling Technology Cat#8054), ULK1 (S757) (Cell Signaling Technology Cat#14202), TMEM106B (Bethyl Labs Cat#A303439A-M), ASAH1 (Proteintech Cat#11274-1-AP), HEXB (Proteintech Cat#16229-1-AP), and BETA-ACTIN (Santa Cruz Biotechnology Cat#69879).

### Molecular cloning

The entry vector pDONR223 containing the full-length GRN (1–1179 base pairs) from the human orfeome collection was used to engineer a stop codon by site-directed mutagenesis (New England Biolabs) at the end of the open-reading frame sequence (ORF). Gateway technology (Thermo Fisher) was used to transfer the GRN ORF using LR cloning from the entry vector to the pHAGE lentiviral destination expression vector. sgRNA sequences for editing the *TMEM192* and *GRN* loci were cloned into the pX459 V2.0 vector (Addgene Cat#62988) as described^31^.

### Gene editing

CRSIPR/Cas9-mediated gene editing of HeLa cells (ATCC Cat# CCL-2) was performed as described^31^. The following sgRNA sequences were used:

TMEM192 (5’-AGTAGAACGTGAGAGGCTCA-3’)

GRN (5’-ATCGACCATAACACAGCACG-3’)

To engineer the lyso-IP tag into HeLa cells by homology-directed repair, a gene block encoding a 3xHA epitope tag, a puromycin cassette, and homology arms on either side of the cleavage site was synthesized by Integrated DNA Technologies to edit the *TMEM192* locus. This sequence was cloned into the pSmart (Lucigen Cat#40041-2) shuttle vector using Gibson assembly (New England Biolabs). The shuttle vector along with the TMEM192 sgRNA sequence was transiently transfected into HeLa cells and puromycin selection was performed 5 days post-transfection for 7–8 days. The mixed pool of cells that were puromycin resistant were single-cell plated, and clonal lines of homozygous HeLa TMEM192^3xHA^ were isolated.

### Lentivirus production

The lentiviral vector was packaged in HEK293T (ATCC Cat#CRL-3216) by cotransfection of psPAX2, pMD2.G (Addgene Cat#12260 Cat#12259) and pHAGE-GRN in a 4:2:1 ratio using polyethelenimine. Virus-containing supernatant was harvested 2 days after transfection and filtered through a 0.45-micron syringe filter. Polybrene was added to a final concentration of 8 μg/ml to the viral supernatant. HeLa^TMEM192 3xHA^ GRN KO cells were infected with 50 μL of viral supernatant and stable cell lines were selected 48 hours post-infection using hygromycin at a concentration of 100 μg/mL.

### Cell culture

HeLa ^TMEM192 3xHA^ and HEK293T cells were grown at 37°C in Dulbecco’ Modified Eagles Medium (DMEM) (Invitrogen Cat#11995-073), supplemented with 10% fetal bovine serum (FBS) (HyClone Cat#SH30910.03) and 1% penicillin-streptomycin (Thermo Fisher Cat#15140163).

### Immunofluorescence and imaging analysis

Cells were plated on to 24-well glass-bottom dish (Cellvis Cat#P24-1.5H-N). All immunofluorescence experiments were performed at room temperature. For the membrane damage experiment, cells were either untreated (DMSO) or treated with 500 μM LLoMe (L-leucyl-L-leucine methyl ester hydrochloride, Cayman Chemical) for 30 minutes. Cells at 70% confluence were washed twice in PBS and fixed with 4% paraformaldehyde in PBS for 20 minutes. Cells were solubilized in 0.25% Triton-X detergent in PBS for 15 minutes and then blocked with 2% BSA in PBS (blocking buffer) for 30 minutes. For the GM2-LAMP1 co-stain experiment, the cells were fixed and blocked, but not treated with detergent. Immunostaining for 2 hours was performed with the following primary antibodies (1:100 dilution in blocking buffer): GM2 (Tokyo Chemical Industry Cat#2576), LAMP1 (Cell Signaling Technology Cat#9091), LGALS3 (Santa Cruz Biotechnology sc23938), and UBIQUITIN (Enzo Life Sciences Cat#BML-PW0930-0100). Cells were washed 3×5 minutes with PBS. Cells were incubated with the appropriate Alexa Fluor-conjugated secondary antibodies (Thermo Fisher) (1:400 dilution in blocking buffer) for 1 hour. Cells were washed 3×5 minutes with PBS. Cells were stained with Hoechst (1 μg/mL) for 5 minutes. Cells were washed 3×5 minutes with PBS. Cells were imaged using a Yokogawa CSU-X1 spinning-disk confocal on a Nikon Ti-E inverted microscope at the Nikon Imaging Center in Harvard Medical School. The microscope is equipped with a Nikon Plan Apo 40x/1.30 NA objective lens and 445 nm (75 mW), 488 nm (100 mW), 561 nm (100 mW) and 642 nm (100 mW) laser lines controlled by AOTF. All images were collected with a Hamamatsu ORCA-ER cooled CCD camera (6.45 μm^2^ photodiode) with MetaMorph image acquisition software. Z series are displayed as maximum z-projections and brightness and contrast were adjusted for each image equally and then converted to RGB. Image analysis was performed using Fiji^32^

### Western blotting analysis

Cell pellets or mouse tissues were resuspended in ice-cold 8 M urea buffer (8 M urea, 50 mM Tris pH 7.4, 50 mM NaCl) supplemented with protease and phosphatase inhibitors (Roche). The resuspended samples were sonicated, and the lysates were clarified at 13,000 rpm for 10 minutes at 4°C. A Bradford assay was performed, and equal amounts of lysate were boiled in LDS-Laemmli buffer supplemented with 50 mM DTT for 10 minutes at 95°C. Lysates were run on 4–20% Tris Glycine gels (BioRad) and transferred on to PVDF membranes (Millipore), which were blocked for 1 hour at room temperature in 2% BSA-0.1% TBS-tween (blocking buffer). Immunoblotting with primary antibodies was performed overnight at 4°C (1:1000 dilution in blocking buffer). Immunoblots were incubated with the appropriate secondary antibodies (1:5000 dilution in blocking buffer) for 1 hour at room temperature. Images of blots were acquired using Enhanced-Chemi luminescence on a BioRad ChemiDoc imager.

### Immunoprecipitation

Cells pellets were harvested, washed twice in ice-cold PBS, and resuspended either in ice-cold neutral lysis buffer (50 mM Tris pH 7.4, 150 mM NaCl, 0.5% NP-40, pH 7.4) or acidic lysis buffer (50 mM NaOAc pH 5.3, 150 mM NaCl, 0.5% NP-40, pH 5.2) supplemented with protease and phosphatase inhibitors (Roche). All subsequent steps were performed at 4°C. Resuspended cells were incubated on a rotator for 30 minutes, centrifuged at 13,000 rpm for 10 minutes, and the supernatant was collected. 1 mg of supernatant from each sample was incubated with 20 μL of BMP-beads, LPA-beads, or naked-beads (Echelon Biosciences Cat#P-BLBP-2 Cat#L-6101 Cat#P-B000) for 3 hours on a rotator. The beads were pelleted by centrifugation at 1000 rpm for 1 minute, the supernatant aspirated, and washed twice with either neutral or acidic lysis buffer. The beads were boiled in LDS-Laemmli buffer supplemented with 50 mM DTT for 10 minutes at 95°C and western blotting was performed as described earlier.

### Lysosome purification

Lysosome immunoprecipitation was carried out as described^33^ with a few modifications. All steps of the process were carried out at 4°C with cold solutions. Briefly, TMEM192^3xHA^ endogenously tagged cells that were grown to 80% confluency in 150-mm plates were washed twice with PBS, scraped into tubes, and then pelleted at *500g* for 5 minutes. The cells were re-suspended in 2 mL of lysoIP buffer (50 mM KCl, 100 mM KH_2_PO_4_, 100 mM K_2_HPO_4_ pH7.2) supplemented with protease and phosphatase inhibitors (Roche). The cells were transferred to a glass homogenizer and dounced using 25 strokes. The lysed cells were centrifuged at 1000*g* for 10 minutes and the post-nuclear supernatant (PNS) was collected. The concentration of the PNS was determined by Bradford assay. Lysosomes from normalized amounts of PNS were immunoprecipitated by incubation with 50 μL of magnetic HA-beads (Thermo Scientific Cat#88837) for 1 hour on a rotator. The beads were sequentially washed once with lysoIP buffer containing 300 mM NaCl and once with lysoIP buffer. The lysosomes were solubilized and eluted off the beads by incubating the beads with 150 μL of lysoIP buffer with 0.5% NP-40 in a thermomixer (1000 rpm) for 30 minutes. The eluates were snap-frozen in liquid nitrogen and stored in −80°C.

### Mice

Animal procedures were approved by the Institutional Animal Care and Use Committee of the Harvard Medical Area Standing Committee on Animals and followed NIH guidelines. Mice were housed in a pathogen-free barrier facility with a 12 h light/12 h dark cycle and allowed food and water ad libitum. *Grn^-/-^* mice^2^ and *Grn^R493X^* mice^12^ were on the C57BL/6J background (backcrossed more than eight generations).

### Human brain tissue studies

Post mortem brain samples were provided by the University of California, San Francisco (UCSF) Neurodegenerative Disease Brain Bank. Brains were donated with the consent of the participants or their surrogates in accordance with the Declaration of Helsinki, and the research was approved by the UCSF Committee on Human Research. Tissue blocks were dissected from the middle frontal gyrus and lateral occipital cortex of three controls, as well as six patients with sporadic FTLD-TDP and thirteen with GRN-FTLD-TDP. All patients with FTD-*GRN* carried a pathogenic variant in *GRN* and had FTLD-TDP, Type A, identified at autopsy (Table see below). Clinical and neuropathological diagnoses were made using standard diagnostic criteria ^34–38^.

| Case P. | Gene | Sex | Age at death | Clinical | Primary Neuropath Dx | PMI (hrs) |
| --- | --- | --- | --- | --- | --- | --- |
| 1 |  | F | 98 | Control | None | 17.0 |
| 2 |  | F | 86 | Control | None | 6.4 |
| 3 |  | M | 76 | Control | None | 8.2 |
| 4 | *GRN* | F | 73 | PNFA/CBS | FTLD-TDP-A | 20.7 |
| 5 | *GRN* | M | 68 | bvFTD | FTLD-TDP-A | 13.5 |
| 6 | *GRN* | F | 70 | PPA, unspecified | FTLD-TDP-A | 9.1 |
| 7 | *GRN* | M | 74 | PNFA/CBS | FTLD-TDP-A | 30.9 |
| 8 | *GRN* | F | 66 | bvFTD | FTLD-TDP-A | 7.4 |
| 9 | *GRN* | M | 64 | bvFTD | FTLD-TDP-A | 7.2 |
| 10 | *GRN* | F | 59 | CBS | FTLD-TDP-A | 9.5 |
| 11 | *GRN* | F | 56 | bvFTD | FTLD-TDP-A | 7.6 |
| 12 | *GRN* | F | 78 | mixed FTD | FTLD-TDP-A | 19 |
| 13 | *GRN* | F | 66 | bvFTD | FTLD-TDP-A | 17.1 |
| 14 | *GRN* | F | 64 | CBS | FTLD-TDP-A | 10.5 |
| 15 | *GRN* | F | 78 | AD vs. FTD | FTLD-TDP-A | 8.1 |
| 16 | *GRN* | M | 72 | AD | AD, FTLD-TDP-A? | 7.2 |
| 17 | Sporadic | M | 72 | PPA-mixed | FTLD-TDP-A | 23.8 |
| 18 | Sporadic | M | 63 | CBS | FTLD-TDP-A | 15.7 |
| 19 | Sporadic | F | 73 | bvFTD | FTLD-TDP-A | 7.4 |
| 20 | Sporadic | F | 78 | PNFA | FTLD-TDP-A | 9.0 |
| 21 | Sporadic | F | 66 | CBS | FTLD-TDP-A | 17.3 |
| 22 | Sporadic | M | 70 | bvFTD | FTLD-TDP-A | 8.5 |

### Lipidomics of the organic phase derived from cells and mouse tissues

HeLa cells were grown in a 10-cm culture dish until they reached ~80% confluence. Tissue samples were obtained as described^12^, mouse brains were collected and immediately snap frozen in liquid nitrogen. Cell and tissue homogenates were obtained by snap-freezing (in liquid nitrogen)/thawing (using an ultrasound water bath for 3 min) repeatedly, and finally extracted according to Folch’s method^39^. Internal standard SPLAH mix and deuterated ganglioside standard spiked in prior to extraction were used for normalization. The organic phase of each cell-culture sample was normalized by total soluble protein amounts and measured by BCA assay (Thermo Scientific, 23225, Waltham, MA), whereas tissue samples were normalized according to dry weight measurements. Samples were routinely subjected to two rounds of extraction. The HPLC-MS method was adopted from^40^. Briefly, HPLC analysis of the organic phases was performed employing a C30 reverse-phase column (Thermo Fisher Scientific, Acclaim C30, 2.1 x 250 mm, 3 μm, operated at 55°C; Bremen, Germany) connected to a Dionex UltiMate 3000 HPLC system and a QExactive orbitrap mass spectrometer (Thermo Fisher Scientific, Bremen, Germany) equipped with a heated electrospray ionization (HESI) probe. Dried lipid samples were dissolved in appropriate volumes of 2:1 MeOH:CHCl3 (v/v), and 5 μl of each sample was injected, with separate injections for positive and negative ionization modes. Mobile phase A consisted of 60:40 ACN/H_2_O, including 10 mM ammonium formate and 0.1% formic acid, and mobile phase B consisted of 90:10 2-propanol/ACN, also including 10 mM ammonium formate and 0.1% formic acid. The elution was performed with a gradient of 90 minute; during 0–7 minutes, elution starts with 40% B and increases to 55%; from 7 to 8 minutes, increase to 65% B; from 8 to 12 minutes, elution is maintained with 65% B; from 12 to 30 minutes, increase to 70% B; from 30 to 31 minutes, increase to 88% B; from 31 to 51 minutes, increase to 95% B; from 51 to 53 minutes, increase to 100% B; during 53 to 73 minutes, 100% B is maintained; from 73 to 73.1 minutes, solvent B was decreased to 40% and then maintained for another 16.9 minutes for column re-equilibration. The flow-rate was set to 0.2 mL/min. The column oven temperature was set to 55°C, and the temperature of the autosampler tray was set to 4°C. The spray voltage was set to 4.2 kV, and the heated capillary and the HESI were held at 320°C and 300°C, respectively. The S-lens RF level was set to 50, and the sheath and auxiliary gas were set to 35 and 3 units, respectively. These conditions were held constant for both positive and negative ionization mode acquisitions. External mass calibration was performed using the standard calibration mixture every 7 days. MS spectra of lipids were acquired in full-scan/data-dependent MS2 mode. For the full-scan acquisition, the resolution was set to 70,000, the AGC target was 1e6, the maximum injection time was 50 msec, and the scan range was m/z = 133.4–2000. For data-dependent MS2, the top 10 ions in each full scan were isolated with a 1.0 Da window, fragmented at a stepped normalized collision energy of 15, 25, and 35 units, and analyzed at a resolution of 17,500 with an AGC target of 2e5 and a maximum injection time of 100 msec. The underfill ratio was set to 0. The selection of the top 10 ions was subject to isotopic exclusion with a dynamic exclusion window of 5.0 sec. Processing of raw data was performed using LipidSearch software (Thermo Fisher Scientific/Mitsui Knowledge Industries)^41,42^.

### Lipidomics of the aequous phase derived from cells and tissues

The aqueous phase was desalted by applying to a C18 cartridge (Waters) equilibrated with 2:43:55 (chloroform-methanol-water) and eluted with 1:1 CHCl3:MeOH. The eluates were dried down again and resuspended in chloroform-methanol-water (600:425:75, v/v/v).

The HILIC-MS method was adopted from^43^. HPLC analysis was performed employing a Phenomenex (Thermo Fisher Scientific, CAT#, 2.0 x 150 mm, operated at 60° C; Bremen, Germany). Dried lipid samples were dissolved in appropriate volumes of 2:1 MeOH: CHCl3 (v/v) and 5μl of each sample was injected and acquired in negative ionization mode. Mobile phase A consisted of acetonitrile with 0.2 %formic acid and mobile phase B consisted of 10Mm aqueous ammonium acetate pH 6.1 adjusted by formic acid. Column equilibration was performed using 12.3 % B for 5 minutes prior to each run. Chromatographic condition: mobile-phase gradient as follows-0 min: 87.7% A + 12.3% B; 15 min: 77.9% A + 22.1% B. The re-equilibration time between runs is 5 mins. The flow rate for the separation was set to 0.6 mL/min. The column oven temperature was set to 40 °C, and the temperature of the autosampler tray was set to 4°C. The spray voltage was set to −4.5 kV, and the heated capillary and the HESI were held at 300 °C and 250 °C, respectively. The S-lens RF level was set to 50, and the sheath and auxiliary gas were set to 40 and 5 units, respectively. These conditions were held constant during the acquisitions. External mass calibration was performed using the standard calibration mixture every 7 days. MS spectra of lipids were acquired in full-scan/data-dependent MS2 mode. For the full-scan acquisition, the resolution was set to 70,000, the AGC target was 1e6, the maximum injection time was 50 msec, and the scan range was m/z = 700-2500 in the negative ion mode. For data-dependent MS2, the top 10 ions in each full scan were isolated with a 1.0 Da window, fragmented at a stepped normalized collision energy of 25, 35, and 50 units, and analyzed at a resolution of 17,500 with an AGC target of 2e5 and a maximum injection time of 100 msec. The underfill ratio was set to 0. The selection of the top 10 ions was subject to isotopic exclusion with a dynamic exclusion window of 5.0 sec. Processing of raw data was performed in Xcalibur™ software (Thermo Fisher Scientific).

### Lipid extraction for mass spectrometry lipidomics of human brain samples

MS-based lipid analysis was performed by Lipotype GmbH (Dresden, Germany) as described^44^. If not indicated otherwise, 500 μg of tissue were used per extraction. Lipids were extracted using a two-step chloroform/methanol procedure^45^. Samples were spiked with internal lipid standard mixture containing: cardiolipin 16:1/15:0/15:0/15:0 (CL, 50 pmol per extraction), ceramide 18:1;2/17:0 (Cer, 30 pmol), diacylglycerol 17:0/17:0 (DAG, 100 pmol), hexosylceramide 18:1;2/12:0 (HexCer, 30 pmol), lyso-phosphatidate 17:0 (LPA, 30 pmol), lyso-phosphatidylcholine 12:0 (LPC, 50 pmol), lyso-phosphatidylethanolamine 17:1 (LPE, 30 pmol), lyso-phosphatidylglycerol 17:1 (LPG, 30 pmol), lyso-phosphatidylinositol 17:1 (LPI, 20 pmol), lyso-phosphatidylserine 17:1 (LPS, 30 pmol), phosphatidate 17:0/17:0 (PA, 50 pmol), phosphatidylcholine 17:0/17:0 (PC, 150 pmol), phosphatidylethanolamine 17:0/17:0 (PE, 75 pmol), phosphatidylglycerol 17:0/17:0 (PG, 50 pmol), phosphatidylinositol 16:0/16:0 (PI, 50 pmol), phosphatidylserine 17:0/17:0 (PS, 100 pmol), cholesterol ester 20:0 (CE, 100 pmol), sphingomyelin 18:1;2/12:0;0 (SM, 50 pmol), triacylglycerol 17:0/17:0/17:0 (TAG, 75 pmol), GM1-D3 18:1;2/18:0;0 (200 pmol), and cholesterol D6 (Chol, 300 pmol). After extraction, the organic phase was transferred to an infusion plate and dried in a speed vacuum concentrator. 1st step dry extract was re-suspended in 7.5 mM ammonium acetate in chloroform/methanol/propanol (1:2:4, V:V:V) and 2nd step dry extract in 33% ethanol solution of methylamine in chloroform/methanol (0.003:5:1; V:V:V). All liquid handling steps were performed using Hamilton Robotics STARlet robotic platform with the Anti Droplet Control feature for organic solvents pipetting.

### MS data acquisition of human brain samples

Samples were analyzed by direct infusion on a QExactive mass spectrometer (Thermo Scientific) equipped with a TriVersa NanoMate ion source (Advion Biosciences). Samples were analyzed in both positive and negative ion modes with a resolution of Rm/z=200=280000 for MS and Rm/z=200=17500 for MS/MS experiments, in a single acquisition. MS/MS was triggered by an inclusion list encompassing corresponding MS mass ranges scanned in 1 Da increments^46^. Both MS and MSMS data were combined to monitor CE, DAG and TAG ions as ammonium adducts; PC, PC O-, as acetate adducts; and CL, PA, PE, PE O-, PG, PI and PS as deprotonated anions. MS only was used to monitor LPA, LPE, LPE O-, LPI, LPS, and GM2 as deprotonated anions; Cer, HexCer, SM, LPC and LPC O-as acetate adducts and cholesterol as ammonium adduct of an acetylated derivative^47^.

### Lipidomic analysis of gangliosides of human brain samples

Ganglioside classes GM1, GD1, GD2, GD3, GT1, GT2, GT3, GQ1 were extracted and analyzed as follows: Ganglioside in the remaining water phase of the two-step chloroform/methanol procedure were subjected to purification using solid phase extraction (Thermo Scientific SOLA SPE plates, 10 mg/2 mL)^48^. The water phase was loaded on columns pre-washed with chloroform/methanol (2:1, V:V), methanol and methanol/water (1:1, V:V); with the input flow through re-applied three times. Then, columns were washed with water and the elution was carried out two times with methanol and one time with chloroform/methanol (1:1, V:V). Washing and elution steps were carried using a vacuum manifold. Pooled eluates were dried in a speed vacuum concentrator and re-suspended in 33% ethanol solution of methylamine in chloroform/methanol (0.003:5:1; V:V:V). Ganglioside extracts were analyzed by direct infusion on a QExactive mass spectrometer (Thermo Scientific) equipped with a TriVersa NanoMate ion source (Advion Biosciences). Samples were analyzed in negative ion modes with a resolution of Rm/z=200 = 140,000; AGC target of 1e6; maximum injection time of 500 ms and 3 microscans.

### Data analysis and post-processing of human brain samples

Data were analyzed with in-house developed lipid identification software, based on LipidXplorer^49,50^. Data post-processing and normalization were performed using an inhouse developed data management system. Only lipid identifications with a signal-to-noise ratio >5, and a signal intensity fivefold higher than in corresponding blank samples were considered for further data analysis. Lipids were normalized to lipid class-specific internal standards. In case of ganglioside classes for which no suitable lipid class-specific internal standards are available, spectral intensities were normalized to the internal standard GM1-D3 18:1;2/18:0;0 and the normalised intensities further normalised to total lipid content (in pmol) of the sample.

Only results with > 1.5fold change and a p-value < 0.05 were selected from the web browser-based data visualization tool (LipotypeZoom) for levels of lipid species as the percentage of the total lipid amount for each sample.

### Enyzme activity assays

Protein estimation of the lysate was done by BCA assay. The activity of β-hexosaminidase and glucosylceramidase β measured by a fluorometric method using 4-Methylumbelliferyl N-acetyl-β-D-glucosaminide (Sigma, Cat#69585) and 4-Methylumbelliferyl β-D-glucopyranoside (Sigma, Cat#M3633) as substrates, respectively. Reaction mixture consisting either of the substrates (5mM), enzyme fraction, and 100 mM citrate buffer, pH 5.0, in a final volume of 200μL were incubated at 45°C. After 1 hour, the reaction was stopped with 2.5 mL of 0.5 M Na_2_CO_3_ buffer, pH 10.4. The 4-methylumbelliferone released was measured in a Tecan Spark multimode microplate reader with excitation and emission set at 350 and 440 nm, respectively.

### Proteomics - Sample digestion

50 μg of protein extracts from cell pellets or mouse tissue were subjected to disulfide bond reduction with 5 mM TCEP (room temperature, 10 minutes) and alkylation with 25 mM chloroacetamide (room temperature, 20 minutes) followed by methanol–chloroform precipitation. For lysosomal samples, disulfide bond reduction and alkylation was performed using 20 μg of extracts, followed by trichloroacetic acid precipitation. Samples were resuspended in 50 μL of 200 mM EPPS, pH 8.5, and digested at 37°C for 2 hours with LysC protease at a 200:1 protein-to-protease ratio. Trypsin was then added at a 100:1 protein-to-protease ratio and the reaction was incubated for 6 hours at 37°C. Tandem mass tag labeling of each sample was performed by adding 10 μL each of the 20 ng/μL stock of TMTpro reagent along with acetonitrile to achieve a final acetonitrile concentration of approximately 30% (v/v). After incubation at room temperature for 1hr, labeling efficiency of a small aliquot was tested, and the reaction was then quenched with hydroxylamine to a final concentration of 0.5% (v/v) for 15 minutes. The TMTpro-labeled samples were pooled together at a 1:1 ratio. The sample was vacuum centrifuged to near dryness, resuspended in 5% formic acid and subjected to C18 solid-phase extraction (SPE) (Sep-Pak, Waters).

### Proteomics - Off-line basic pH reversed-phase (BPRP) fractionation

Dried TMTpro-labeled sample was resuspended in 100 μl of 10 mM NH_4_HCO_3_, pH 8.0, and fractionated using BPRP HPLC. Briefly, samples were offline fractionated over 90 minutes and separated into 96 fractions by high pH reverse-phase HPLC (Agilent LC1260) through an Agilent ZORBAX 300Extend C18 column (3.5-μm particles, 4.6 mm ID and 250 mm in length) with mobile phase A containing 5% acetonitrile and 10 mM NH_4_HCO_3_ in LC-MS grade H_2_O, and mobile phase B containing 90% acetonitrile and 10 mM NH_4_HCO_3_ in LC-MS grade H_2_O (both pH 8.0). The 96 resulting fractions were then pooled in a non-continuous manner into 24 fractions and 12 fractions (non-adjacent) were used for subsequent mass spectrometry analysis. For lysosomal extracts, the dried TMTpro-labeled sample was resuspended in 300 μL of 0.1% trifluoroacetic acid and then fractionated into six fractions, using the High pH reversed-phase peptide fractionation kit (Thermo Fisher Cat#84868). Fractions were vacuum centrifuged to near dryness. Each consolidated fraction was desalted via StageTip, dried again via vacuum centrifugation, and reconstituted in 5% acetonitrile, 1% formic acid for LC-MS/MS processing.

### Proteomics - Liquid chromatography and tandem mass spectrometry

MS data were collected using an Orbitrap Fusion Lumos mass spectrometer (Thermo Fisher) coupled to a Proxeon EASY-nLC1200 liquid chromatography (LC) pump (Thermo Fisher). Peptides were separated on a 100 μm inner diameter microcapillary column packed in house with ~35 cm of Accucore150 resin (2.6 μm, 150 Å, Thermo Fisher) with a gradient consisting of 5–21% (0–125 min), 21–28% (125–140 min) (ACN, 0.1% FA) over a total 150 min run at ~500 nL/min. For analysis, we loaded 2-3 μg of each fraction onto the column. Each analysis used the Multi-Notch MS^3^-based TMT method, to reduce ion interference compared to MS^2^ quantification. The scan sequence began with an MS^1^ spectrum (Orbitrap analysis; resolution 120,000 at 200 Th; mass range 400-1400 m/z; automatic gain control (AGC) target 5×10^5^; maximum injection time 50 ms). Precursors for MS^2^ analysis were selected using a Top10 method. MS^2^ analysis consisted of collision-induced dissociation (quadrupole ion trap analysis; Turbo scan rate; AGC 2.0×10^4^; isolation window 0.7 Th; normalized collision energy (NCE) 35; maximum injection time 35 ms). Monoisotopic peak assignment was used, and previously interrogated precursors were excluded using a dynamic window (120 s ±10 ppm). After acquisition of each MS^2^ spectrum, a synchronous-precursor-selection (SPS) MS^3^ scan was collected on the top 10 most intense ions in the MS^2^ spectrum^34^. MS^3^ precursors were fragmented by high energy collision-induced dissociation (HCD) and analyzed using the Orbitrap (NCE 55; AGC 3×10^5^; maximum injection time 100 ms, resolution was 50,000 at 200 Th).

### Proteomics-Data analysis

Raw mass spectra obtained were processed using Sequest. Mass spectra was converted to mzXML using a version of ReAdW.exe. Database searching included all entries from the Human Reference Proteome. Searches were performed with the following settings 1) 20 ppm precursor ion tolerance for total protein level analysis, 2) Product ion tolerance was set at 0.9 Da, 3) TMTpro on lysine residues or N-termini at +304.207 Da 4) Carbamidomethylation of cysteine residues (+57.021 Da) as a static modification and oxidation of methionine residues (+15.995 Da) as a variable modification. Peptide-spectrum matches (PSMs) were adjusted to a 1% false discovery rate (FDR)^51^ PSM filtering was performed using a linear discriminant analysis. To quantify the TMTpro-based reporter ions in the datasets, the summed signal-to-noise (S:N) ratio for each TMTpro channel was obtained and found the closest matching centroid to the expected mass of the TMTpro reporter ion (integration tolerance of 0.003 Da). Proteins were quantified by summing reporter ion counts across all matching PSMs, as described previously^52^ PSMs with poor quality, or isolation specificity less than 0.7, or with TMTpro reporter summed signal-to-noise ratio that were less than 100 or had no MS^3^ spectra were excluded from quantification. Values for protein quantification were exported and processed using Perseus to calculate Log fold-changes and p-values. Volcano plots using these values were plotted in Excel.

### Thin-layer chromatography analysis

Analytic thin-layer chromatography (TLC) was performed on 10 cm high-performance thin-layer chromatography (HPTLC) plates (Sigma, Cat# 1056310001). Organic fraction of samples were dried down and analyzed by 2D TLC with chloroform-methanol-water-concentrated ammonia 70:30:3:2 (by vol) used as the first dimension and chloroform-methanol-water 65:35:5 (by vol) used as the second dimension as described previousl^53^. Standard lipids (10 μg) dissolved in methanol or chloroform-methanol (1:1, v/v) were used as a reference.

### Statistical analysis

All statistical analysis was performed using GraphPad Prism 8. Information about significance test is provided in the respective figure legends. All multiple comparisons were performed with the Dunn multiple comparisons correction. Statistically significant differences are denoted as follows: *p<0.05, **p<0.01, ***p<0.001, #p<0.0001.

